# Autophagy mediates temporary reprogramming and dedifferentiation in plant somatic cells

**DOI:** 10.1101/747410

**Authors:** Eleazar Rodriguez, Jonathan Chevalier, Jakob Olsen, Jeppe Ansbøl, Vaitsa Kapousidou, Zhangli Zuo, Steingrim Svenning, Christian Loefke, Stefanie Koemeda, Pedro Serrano Drozdowskyj, Jakub Jez, Gerhard Durnberger, Fabian Kuenzl, Michael Schutzbier, Karl Mechtler, Signe Lolle, Yasin Dagdas, Morten Petersen

## Abstract

Somatic cells acclimate to changes in the environment by temporary reprogramming. Much has been learned about transcription factors that induce these cell-state switches in both plants and animals, but how cells rapidly modulate their proteome remains elusive. Here, we show rapid induction of autophagy during temporary reprogramming in plants triggered by phytohormones, immune and danger signals. Quantitative proteomics following sequential reprogramming revealed that autophagy is required for timely decay of previous cellular states and for tweaking the proteome to acclimate to the new conditions. Signatures of previous cellular programs thus persist in autophagy deficient cells, affecting cellular decision-making. Concordantly, autophagy deficient cells fail to acclimatize to dynamic climate changes. Similarly, they have defects in dedifferentiating into pluripotent stem cells, and redifferentiation during organogenesis. These observations indicate that autophagy mediates cell state switches that underlie somatic cell reprogramming in plants and possibly other organisms, and thereby promotes phenotypic plasticity.

## Introduction

Somatic cells in multicellular eukaryotes are relentlessly exposed to diverse physiological and environmental stimuli including changes in temperature, nutrients, hormones and pathogen load (Cherkasov *et al*, 2013; Chovatiya & Medzhitov, 2014). At certain levels such stimuli become stressful and provoke adaptive cellular responses (Galluzzi *et al*, 2018). To survive, eukaryotes have evolved sophisticated acclimation mechanisms that mediate temporal reprogramming of somatic cells. In both animals and plants, these mechanisms include alterations in transcriptional activities and epigenetic signatures (Davière & Achard, 2016; Hafner *et al*, 2019; Koo & Guan, 2018; Xu *et al*, 2017; Zhang *et al*, 2018).

Somatic cells can also undergo directional reprogramming through dedifferentiation and can form pluripotent cells. This allows somatic cells to redifferentiate into other cell types, organs and even whole organisms in plants (Ikeuchi *et al*, 2015; Li & Belmonte, 2017; Papp & Plath, 2013; Li *et al*, 2017; Takahashi & Yamanaka, 2006). Similar to temporary reprogramming, reprogramming into other cell types is orchestrated by evolutionarily conserved processes and involves major changes in the transcriptome and epigenetic landscape (Iwafuchi-Doi, 2019; Roche *et al*, 2017; Sang *et al*, 2018). Despite the wealth of knowledge on initial transcriptional and epigenetic changes driving somatic reprogramming events, how proteostasis mechanisms delete current cellular states to allow new programs to be installed remain largely unknown.

Macroautophagy (hereafter autophagy) is a conserved quality-control pathway that facilitates cellular adaptation by removing superfluous or damaged macromolecules and organelles (Ho *et al*, 2017; Liu & Klionsky, 2015; Popovic & Dikic, 2014). Although initially discovered as a starvation induced survival mechanism in yeast (Yang & Klionsky, 2013), many studies have now shown that it plays crucial roles in a variety of stress responses (Dikic & Elazar, 2018; Katheder *et al*, 2017; Kumsta *et al*, 2017; Mizushima *et al*, 2008; Munch *et al*, 2014; Bassham *et al*, 2006; Rui *et al*, 2015), and may act both as positive and negative regulator of programmed cell death (Berry & Baehrecke, 2007; Gutierrez *et al*, 2004; Hofius *et al*, 2009; Liu *et al*, 2005; Nakagawa *et al*, 2004). Similarly, autophagy has also been implicated in induced pluripotent stem cell formation, cellular regeneration and stem cell survival in mammals (Boya *et al*, 2018; Calvo-Garrido *et al*, 2019; Saera-Vila *et al*, 2016). However, some of these studies have contrasting conclusions. For example, autophagy was shown to have opposite functions in mammalian cells during reprogramming into pluripotency (Wang *et al*, 2013; Wu *et al*, 2015) and stem cell maintenance in mice (Ho *et al*, 2017; Mortensen *et al*, 2011).

So, how can we reconcile these functions and discrepancies regarding the function of autophagy? And is autophagy involved in induced pluripotent stem cell formation in plants? Unlike reprogramming in stem cells, temporary reprogramming events in somatic cells are reversible and provide phenotypic plasticity in response to various stimuli (Fusco & Minelli, 2010; Kelly *et al*, 2012; Oostra *et al*, 2018; Pfennig *et al*, 2010). Autophagy possesses the degratory capacity to rapidly attenuate cellular programs, and to allow new programs to unfold while also modulating their intensity. Autophagy may therefore be engaged to adjust cell state switching in response to different stimuli.

Here, we find that autophagy functions in many types of cellular reprogramming in plants. Stimuli as diverse as phytohormones, danger signals and microbial elicitors all trigger rapid and robust activation of autophagy. Using quantitative proteomics, we show that autophagy mediates the switch between somatic cell programs by removing cellular components that are no longer required. At the same time, autophagic mechanisms ensure a controlled execution of the newly established programs. Accordingly, autophagic dysfunction leads to defects in organismal fitness, dedifferentiation of somatic cells into pluripotency, and redifferentiation of pluripotent cells into other cell types in plants.

## Results

### Autophagy is rapidly engaged upon perception of diverse stimuli

To examine if temporary reprogramming engages autophagy, we exposed young seedlings of the model plant *Arabidopsis thaliana* expressing the autophagic markers GFP-ATG8a or YFP-mCherry NBR1 (Svenning *et al*, 2011) to an array of treatments. We evaluated autophagic flux in response to a selection of microbial elicitors, danger signals and hormones known to induce temporary reprogramming:Peptide-1 (PEP1, small peptide produced during wounding) and ATP, which are perceived as danger associated molecular patterns (DAMP); Abscisic Acid (ABA, a hormone commonly associated with abiotic stress responses); ACC (1-aminocyclopropane-1-carboxylic acid, precursor of the gaseous hormone ethylene involved in development and senescence); Brassinolide (BL, a steroid hormone involved in growth); NAA (1-Naphthalene Acetic Acid, an auxin involved in growth modulation); and 6-BA (a cytokinin involved in cytokinesis and growth). These treatments induced rapid accumulation of GFP-ATG8a (Fig. 1a, b) and YFP-mCherry NBR1 foci (Supplementary Fig. 1a, b). GFP-ATG8a vacuolar degradation produces free GFP fragments that can be detected by immunoblotting to measure autophagic flux (Mizushima 2010). All of the treatments induced accumulation of free GFP, pointing to increased autophagic flux (Fig. 1c). Further corroboration of autophagic flux increase came from immunoblotting against native NBR1, a well-known autophagy receptor (Fig. 1d). Because high NBR1 turnover complicates this analysis in wild type plants, we used *atg2-2* (Wang *et al*, 2011) instead and observed increased levels of NBR1 in all treatments (Fig. 1d), further confirming the induction of autophagy during temporary reprogramming events. Taken together, our results indicate that regardless of the nature of the signal, autophagy is rapidly induced and may function as an intrinsic component in temporary reprogramming of somatic cells.

**Fig. 1.**
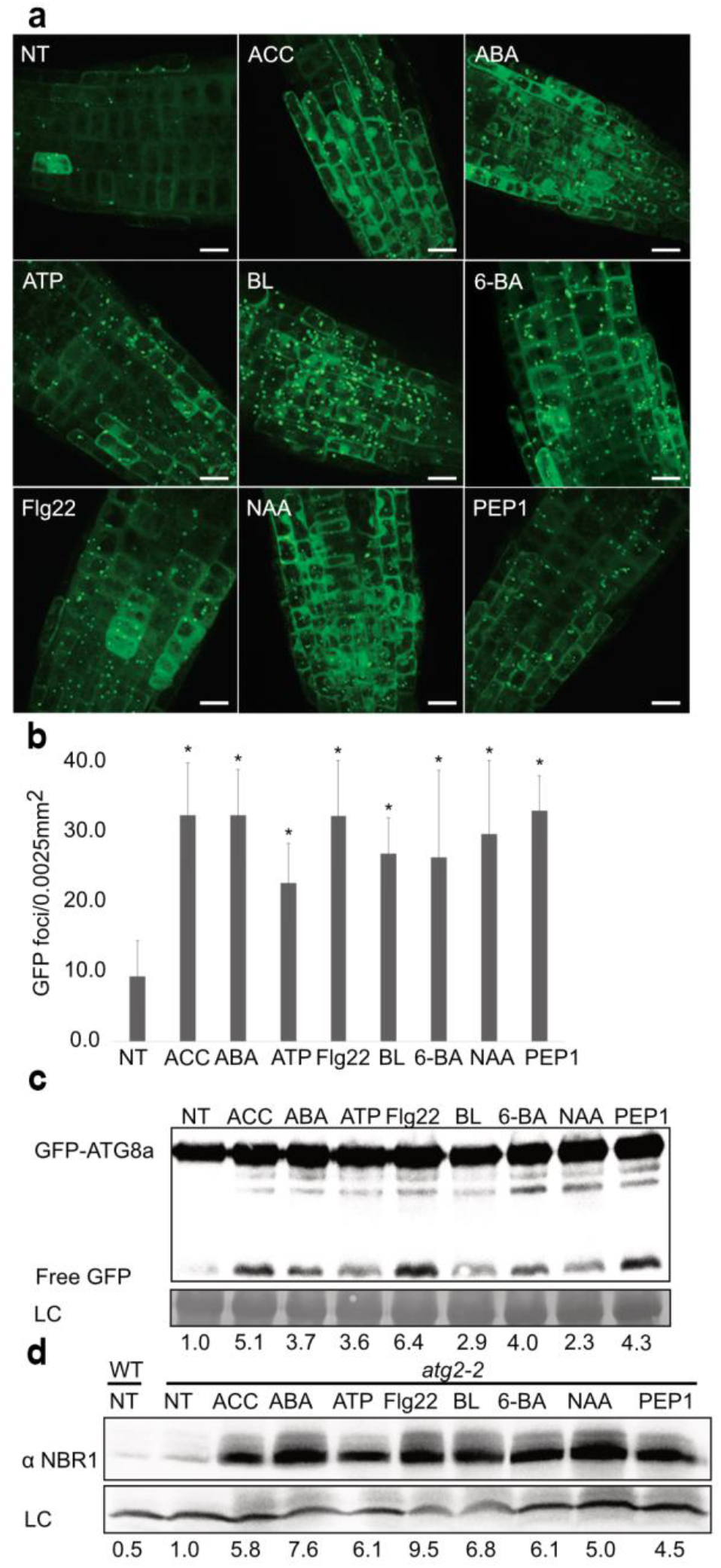
Autophagy is rapidly engaged upon perception of diverse stimuli. GFP-ATG8a expressing seedlings in Murashige & Skoog (MS) growth medium or 30 min after treatment with MS containing ACC, ABA, ATP, BL, 6-BA, Flg22, NAA or PEP1. (**a**) Representative maximum intensity projection images of 10 Z-stacks per image. Scale bar: 10 µm. (**b**) Quantification of GFP foci per 0.0025 mm^2^. Values were calculated from at least three independent experiments with 3 individuals per replicate. Bars marked with an asterisk (*) are statistically significant (P<0.05). (**c**) GFP-ATG8a cleavage immunoblot for plants exposed to the same treatments as in (**a**). (**d**) NBR1 immunoblot for *atg2-2* samples for given treatments. Numbers below the blots represent ratio for given sample normalized to input and relative to non-treated control. Experiments were repeated minimum 3 times with similar results.

### Autophagy facilitates temporary reprogramming

We hypothesized that the primary function of autophagy in temporary reprogramming is to assist cellular “clean-up” to allow a new program to unfold before returning to basal levels. If so, *(i)* a second reprogramming stimulus should re-activate autophagy, and *(ii)* establishment of the second program should be concurrent with a rapid decay of the first program. To test this, we applied consecutive stimuli and examined reprogramming from ABA (abiotic stress proxy) to flg22 (immunity stress proxy) (Fig. 2a, Supplementary Fig.2A). We quantified GFP-ATG8a foci (Fig. 2a, b), YFP-mCherry-NBR1 (Supplementary Fig 2a, b), and free GFP via the cleavage assay (Fig. 2c). All of these assays demonstrated that autophagic flux resets to basal levels after 16h of ABA treatment, contrasting with the rapid induction seen before (Fig. 1). Transferring those seedlings to flg22-containing medium caused reactivation of autophagy as demonstrated by significant accumulation of GFP and YFP-mCherry-positive foci (P<0.05, Fig. 2b and Supplementary Fig 2b) and free GFP (Fig. 2c), in comparison to the control treatment. Hence, our data indicates that autophagy is engaged to clean up, is reset after the clean up, and can be reactivated upon perception of new stimuli.

**Fig. 2.**
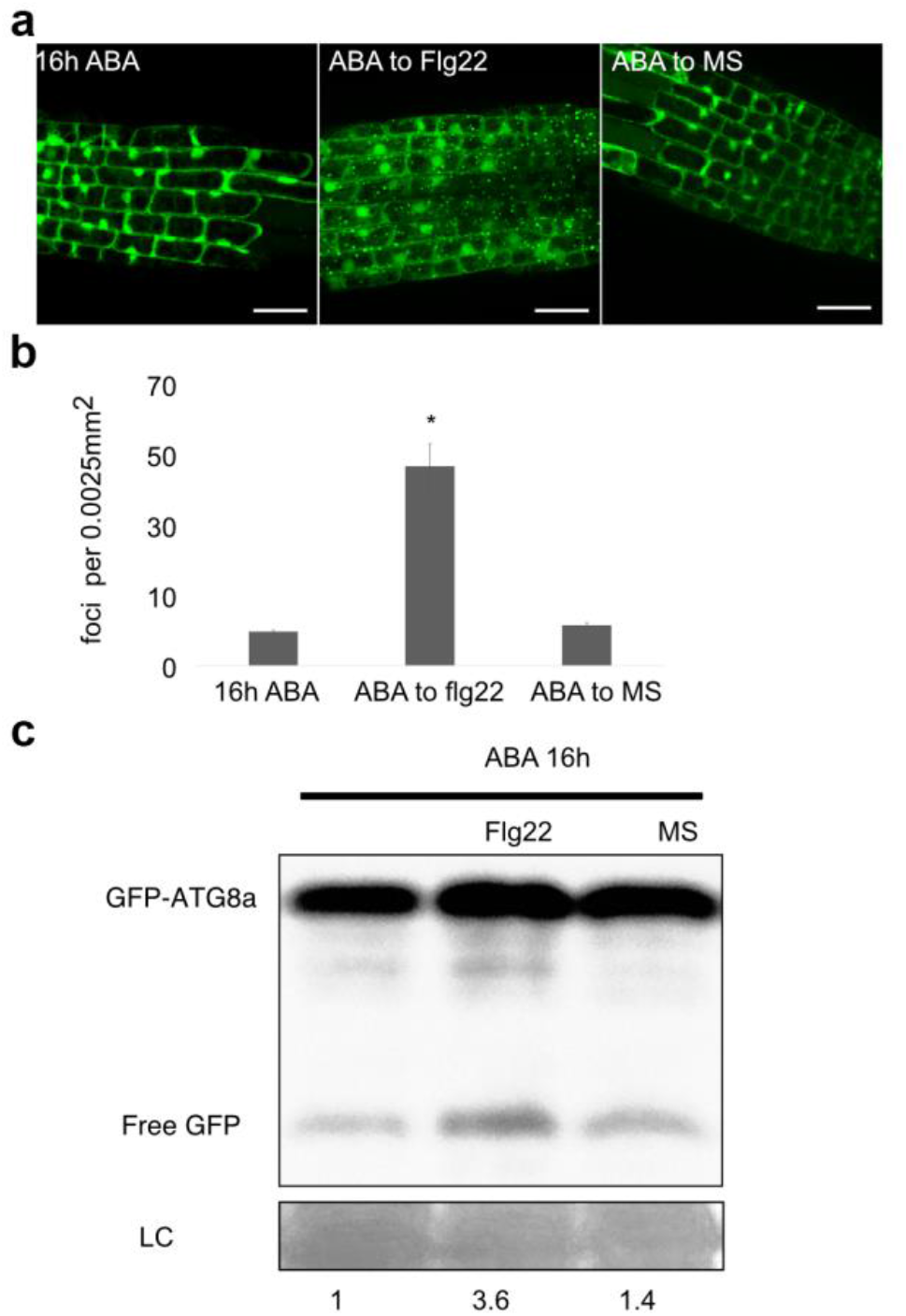
Autophagy is reactivated upon contrasting stimuli perception. Seedlings were acclimated for 16 h in MS containing ABA and then imaged 30 min after being swapped to MS or MS containing flg22. (**a**) Representative maximum intensity projection of 10 Z stacks per image. Experiments were repeated 3 times independently with similar results Scale bar: 10 µm. (**b**) Quantification of GFP foci for given treatments per 0.0025 mm^2^. Values are based on 3 independent experiments, with 3 individuals per condition. Bars marked with an asterisk (*) are statistically significant (P<0.05). (**c**) GFP-ATG8a cleavage immunoblot for samples in **(a)**.

To further support our observations, we performed comparative proteomics using Tandem Mass Tag labelling (TMT) Mass Spectrometry (MS/MS) on wild type (WT) and the autophagy-deficient mutant *atg2-2* upon consecutive, temporary reprogramming stimuli (Fig. 3a). We detected 11,300 proteins, of which 1,241 responded to the treatments (Supplementary Table 1). Based on their behavior, we divided these proteins into fifty clusters. Validating our proteome profiling approach, various ATG8 isoforms and NBR1 clustered together and accumulated to higher levels in *atg2-2* (Supplementary Fig 3a). We then searched for proteins induced by ABA that decreased when switched to flg22 treatment in WT plants but failed to decrease in *atg2-2* (Fig. 3b, correlation = 0.86, 10.5% of all responding proteins). Importantly, most proteins with this profile also decreased faster in WT plants when switched from ABA to flg22 than to control media (Fig. 3c). Several proteins fitting this profile have been previously associated with ABA responses, among them TSPO, which is degraded through autophagy upon completion of the ABA program (Guillaumot *et al*, 2009). Using a TSPO antibody, we confirmed that TSPO follows the same pattern in another autophagy deficient mutant, *atg5-1* (Thompson *et al*, 2005), during consecutive reprogramming from ABA to flg22 (Supplementary Fig. 3b), as well as during reprogramming from ABA to NAA (Supplementary Fig. 3c). These results indicate that autophagy is activated to rapidly remove components of previous cellular programs.

**Fig. 3.**
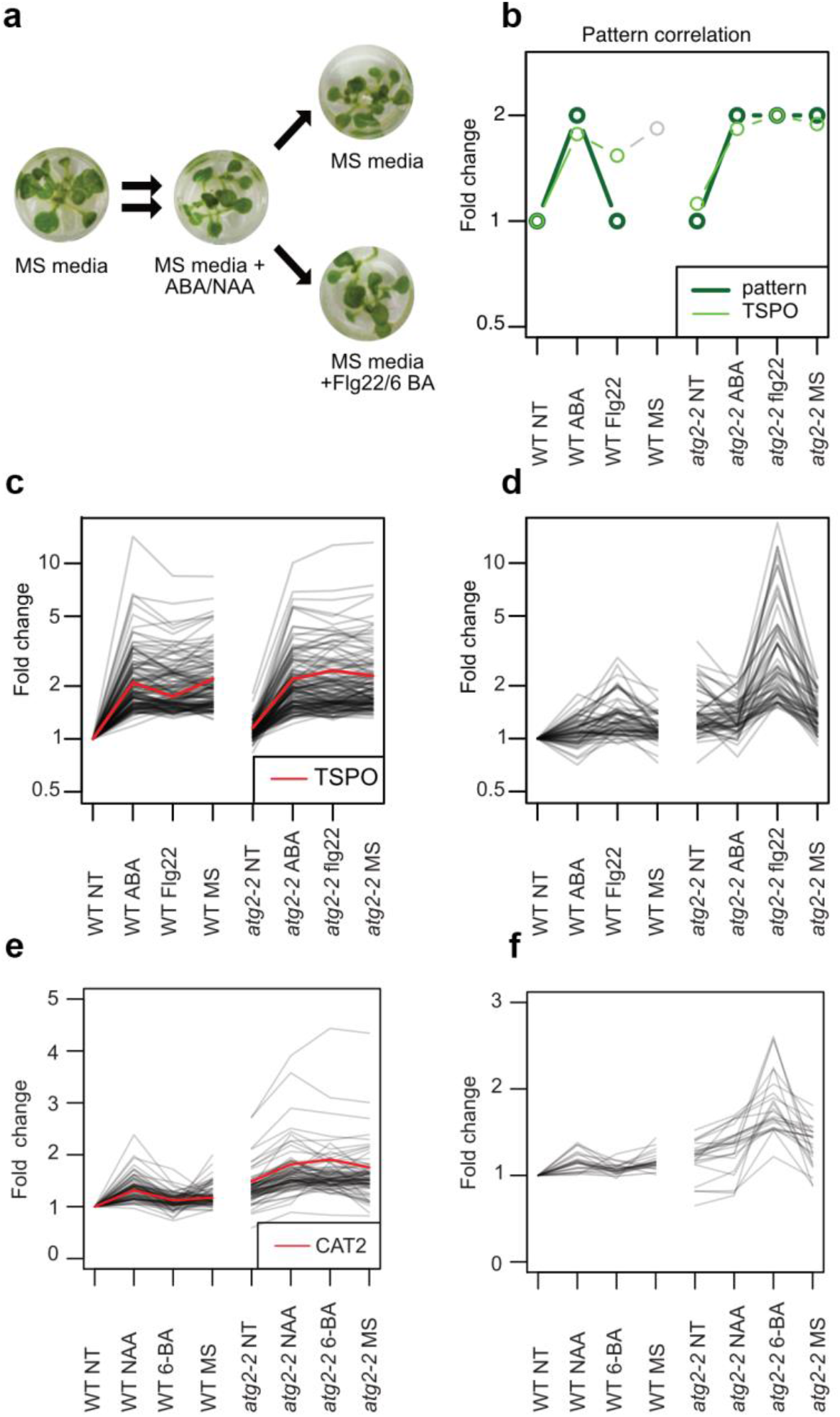
Autophagy facilitates temporary reprogramming during perception of contrasting stimuli by removing old components and modulating the intensity of new responses. (**a**) Schematic representation of the strategy used for consecutive stress treatment. (**b**) Pattern correlation used to find proteins that accumulate upon ABA treatment and are removed in WT but not in *atg2* after swapping to flg22. (**c-e**) Protein clusters obtained after quantitative proteomics of WT and *atg2-2* samples treated as described in (**a**). (**c**) Protein cluster fitting the pattern displayed in (**b**). (**d**) Protein cluster for proteins that accumulate to higher levels in *atg2* than WT upon treatment with flg22. (**e**) Protein cluster of proteins which accumulate upon NAA treatment and are removed in WT but not in atg2 after swapping to 6-BA (**f**) Protein cluster for proteins that accumulate to higher levels in *atg2* than WT upon treatment with 6-BA.

Our clustering analyses revealed that some stress-related proteins peaked to much higher levels in *atg2-2* upon switching from ABA to flg22 (Fig. 3d, 4.6% of all responding proteins). As expected, many of these such as ATPXG2, PDF2.1, and its close homolog AT1G47540 have previously been associated with immune responses (Petersen *et al*, 2000; Tsiatsiani *et al*, 2013; Zhao, 2015). This indicates that autophagy also modulates the intensity of a new cellular program when it is being installed.

To confirm that autophagy functions as an intrinsic component in cellular reprogramming, we extended our proteomics analysis and examined reprogramming between the contrasting developmental phytohormones auxin and cytokinin (Supplementary Table 2). Here we also observed major proteostatic dysregulation in *atg2-2* plants and identified a major cluster (Fig. 3e, 10.2% of all responding proteins), comparable to our previous observations (Fig. 3a). We identified catalase 2 (CAT2) in this cluster and used a catalase antibody to confirm the same pattern in both *atg5-1* and *atg2-2* (Supplementary Fig. 3d). Interestingly, we observed that auxin responsive proteins accumulate in untreated *atg2-2* (Fig. 3e), unlike ABA responsive proteins (Fig. 3a). Since stress programs (ABA) are normally ‘off’ under normal growth conditions, while growth and development programs (auxin) are recruited continuously, gradual accumulation of auxin responsive proteins may not be surprising in autophagy-deficient backgrounds. Similar to our observations above (Fig. 3d), these results again show that autophagy is also needed to modulate the intensity of new cellular programs when switched from auxin to cytokinin (Fig. 3f, 3% of all responding proteins).

### Autophagy deficiencies lead to reduced phenotipic plasticity and increased heterogeneity

The above results indicate that cells lacking autophagic activity lose cellular homeostasis and accumulate signatures of different cellular programs and states. If so, autophagy deficiency may lead to increased heterogeneity during acclimatization to fluctuating environmental conditions. To assess this at an organismal level, we used a high-throughput phenotyping chamber to compare the development of WT, *atg2-2* and *atg5-1* mutant plants grown in standard, stable conditions versus plants grown in highly variable conditions recorded for the Swedish spring of 2013 (Fig. 4a). Data dispersion for dry weight was higher in *atg* plants than in WT, regardless of the conditions tested, albeit with more outliers for *atg2* grown under variable conditions (Fig. 4b, c, Supplementary Fig. 4). This indicates that the loss of cellular homeostasis in *atg* mutants translates into higher heterogeneity, and this increased heterogeneity might stem from their decreased ability to cope with daily variations in temperature, humidity and light conditions that all involve dynamic, temporary reprogramming events.

**Fig. 4.**
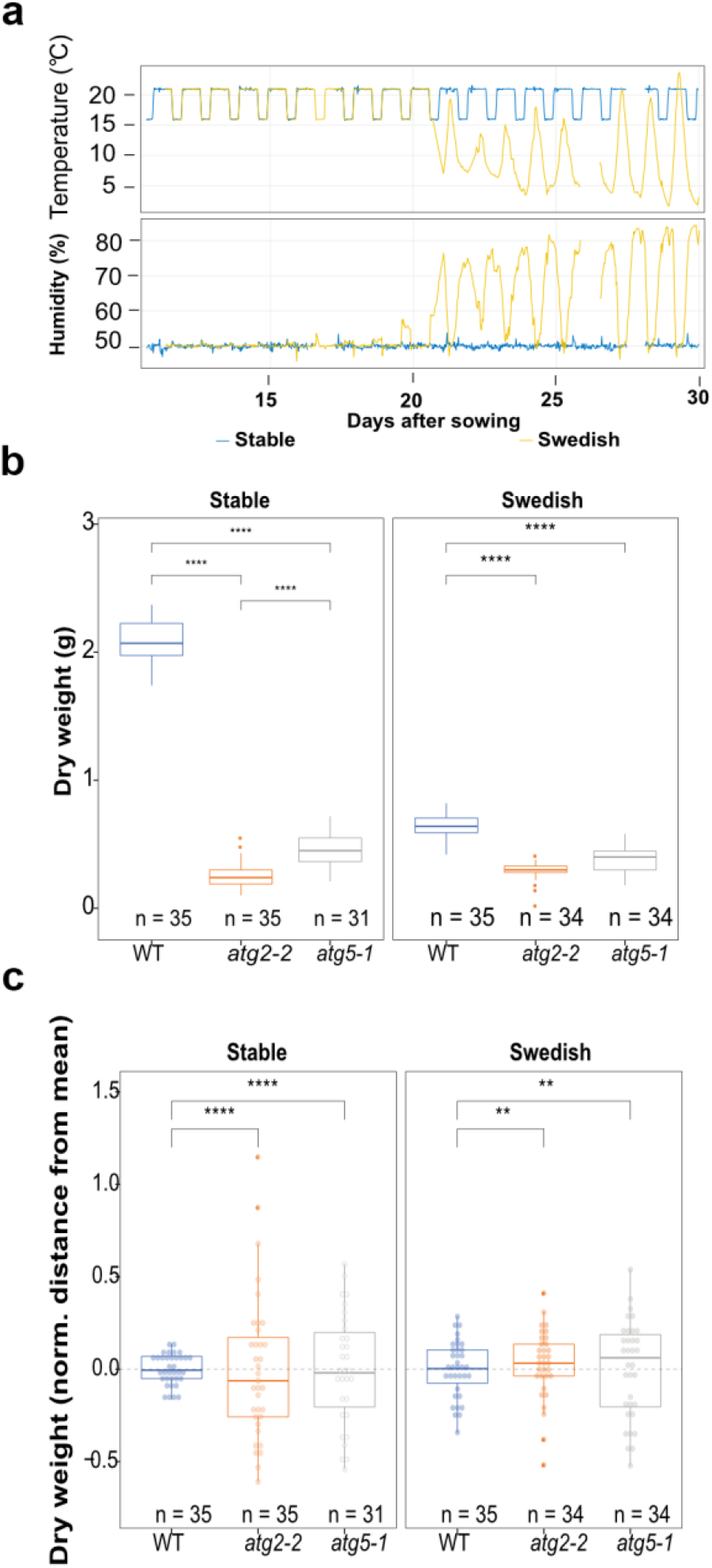
Autophagy deficiencies lead to reduced phenotipic plasticity and increased heterogeneity. (**a**) Weather pattern recorded for the Swedish spring of 2013. **(b)** Dry weight of WT, *atg5-1* or *atg2-2* plants grown under stable (21/16 °C 16/8-h photoperiod) or following the Swedish Spring 2013. **(c)** Box plots displaying the heterogeneity of samples in **(d)** Significance calculated by Kruskal-Wallis or Feltz and Miller test for the equality of coefficients of variation as explained in materials and methods.

### Dedifferentiation is severely impaired in autophagy deficient cells

To substantiate the importance of autophagy in preserving phenotypic plasticity, we examined induced pluripotent stem cell (iPSC) formation and organ regeneration − well-known examples of somatic cell reprogramming − in autophagy deficient plants. Plant pluripotent stem cells (PSCs) can be induced *in vitro* by adjusting the ratios of auxin and cytokinin to trigger dedifferentiation and proliferation of an unorganized mass of pluripotent cells (callus) (Sang *et al*, 2018; Sugimoto *et al*, 2010). When we placed WT, *atg2-2* and *atg5-1* root explants in callus-inducing medium (CIM) to induce PSC and callus formation ((Valvekens *et al*, 1988)), WT root explants dedifferentiated into PSC and formed visible calli, but *atg2* and *atg5* root explants did not (Fig. 5a). When we then transferred these to shoot-inducing medium (SIM) (Valvekens *et al*, 1988) to trigger *de novo* organogenesis, 80% of WT explants formed shoots after 21 days but *atg2-2* (5%) and *atg5* (20%) were severely defective in shoot formation (Fig. 5a, b). These results are consistent with data from zebrafish in which autophagy deficiency impairs the reprogramming that is necessary for muscle regeneration (Saera-Vila *et al*, 2016), and also reduces self renewal and proliferative ability of haematopoetic stem cells (Ho *et al*, 2017).

**Fig. 5.**
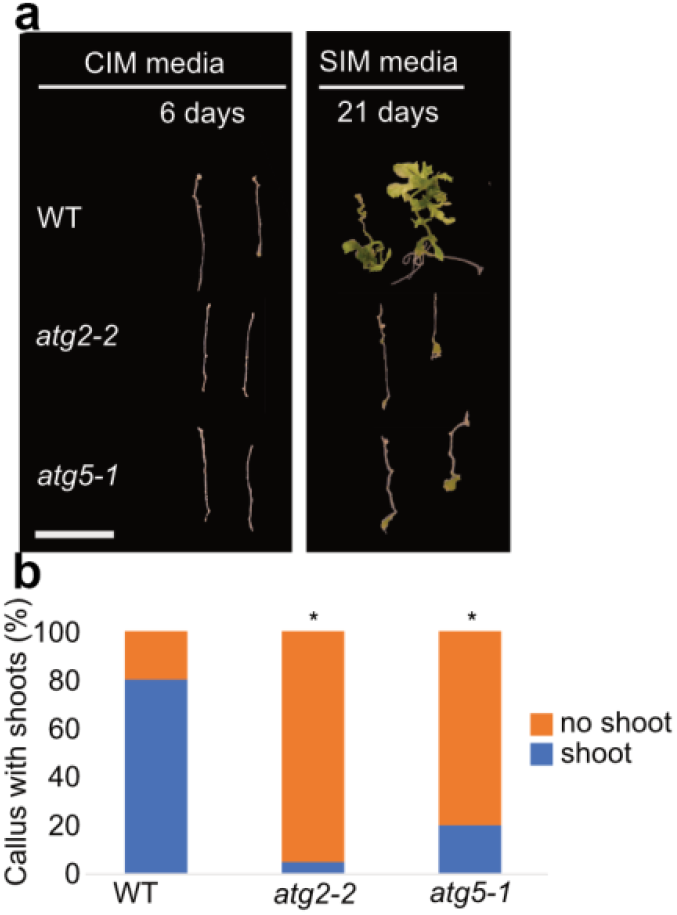
Dedifferentiation is severely impaired in autophagy deficient cells. (**a**) Representative images of root explants from WT, *atg2-2* and *atg5-1* in CIM (6 days) or CIM + SIM (6 + 21 days). (**b**) Quantification (%) of explants presenting shoots after 21 days after incubation on SIM. Results were obtained from 3 independent experiments with at least 20 calli per condition. Bars marked with an asterisk (*) are statistically significant (P<0.05).

Unlike mammals and flowering plants, bryophytes can naturally form PSC without the need for exogenous hormone treatment or overexpression of transcription factors. In particular, the moss *Physcomitrella patens* is able to dedifferentiate somatic cells into chloronema stem cells to repair damaged tissue upon wounding (Ishikawa *et al*, 2011; Kofuji & Hasebe, 2014; Li *et al*, 2017). Using this system, we compared reprogramming efficiency in WT, *atg5* and *atg7* lines upon wounding. As it can be seen in Supplementary Fig. 5a-d, autophagy deficient moss lines are slower and less efficient than WT. These results indicate that, like iPSC formation in *Arabidopsis* (Fig. 5) and zebra fish muscle regeneration (Saera-Vila *et al*, 2016), autophagy deficiency in *P. patens* severely impairs iPSC formation and tissue regeneration. This suggests autophagy mediated dedifferentiation and organogenesis are evolutionary conserved across kingdoms.

### Autophagy deficient cells lose control over *de novo* organogenesis upon prolonged cultivation in pluripotent cell inducing media

Since *atg* mutants made small amounts of callus, we wondered if prolonging CIM treatment could help callus formation in *atg* mutants. We therefore extended the time explants were kept on CIM from 6 to 21 days, while mantaining the subsequent SIM step to 21 days (Protocols, 2009). As expected, at 21 days on CIM, *atg* mutants produced more calli, but still less than WT (Fig. 6a, b). Interestingly, when then moved to SIM, *atg* mutants quickly started to catch up in callus mass (Fig. 6a, c), and produced significantly more callus tissue and shoots (P<0.05) than WT. This is consistent with our proteomics data; since *atg* mutants cannot modulate cellular programs and thus end up with exaggerated calli and shoot formation. This is analogous to comparing acceleration down a developmental ‘slope’ with (WT) or without (*atg* mutants) brakes. These results are in agreement with those of (Ho *et al*, 2017) who desmonstrated that *atg* deficient haematopoietic cells lost their stemmnes and accelerated differentiation. While Ho and colleagues concluded that the enhanced metabolic rate of *atg* deficient cells led to enhanced differentiation, our data also supports a model in which gradual accumulation of conflicting programs, which are not “cleaned” from the cell, could reduce stem cell control.

**Fig. 6.**
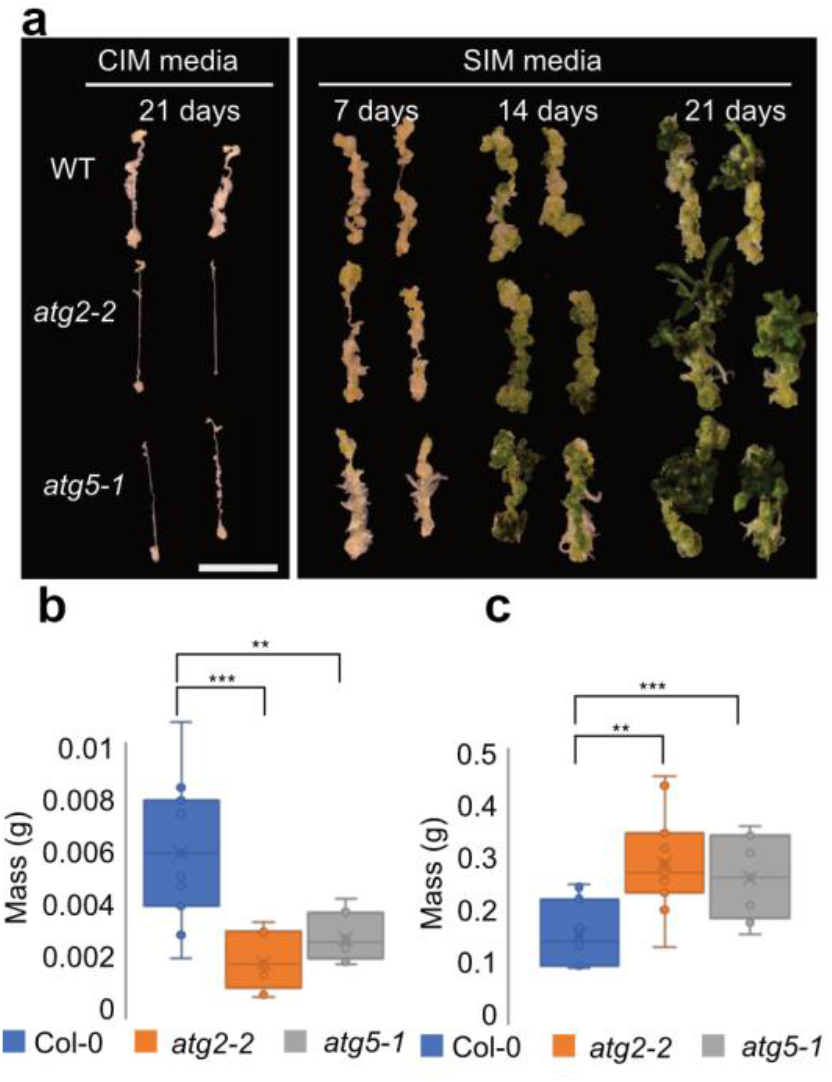
Autophagy deficient cells lose control over *de novo* organogenesis upon prolonged cultivation in pluripotent cell inducing media. (**a**) Representative images of root explants from WT, *atg2-2* and *atg5-1* in CIM (21 days) or CIM + SIM (21+ 21 days). (**b and c**) Fresh weights of calli after incubation on CIM (**b**) or CIM + SIM (**c**). Results were obtained from 3 independent experiments with at least 20 calli per condition, asteriscs mark statistical significance to WT according to the T -test (** P<0.001; *** P<0.0001).

As autophagy helps to fight aging (Fernández *et al*, 2018; Kaushik & Cuervo, 2015; Leidal *et al*, 2018) and preserve mammalian stem cell function (Ho *et al*, 2017; Vilchez *et al*, 2014), we wanted to address this in the context of plant stem cells. For this, we prolongued calli culture on SIM to 5 weeks and observed that both *atg* mutant calli displayed premature senescence and death (Supplementary Fig. 6). Taken together, our results demonstrate that autophagy mediates cellular reprogramming necessary for iPSC formation, modulates subsequent *de novo* organogenesis, and assists in maintaining longevity of plant iPSC masses.

## Conclusion

In summary, our results at the cellular and organismal level show that autophagy is rapidly activated upon perception of diverse stimuli to maintain cellular competence by mediating cellular clean-up during various reprogramming events in plants. Defects in autophagy led to increased heterogeneity and chaotic cellular decisions, presumably because signatures of previous programs are not removed efficiently and interferes with the execution of new programs. Because accumulation of these programs happens gradually during development, the longer cells experience this proteostasis deficiency, the higher becomes the probability for conflicting cellular decisions. Thus, the numerous, apparently opposite conclusions on autophagic functions in different developmental systems (Ho *et al*, 2017; Hofius *et al*, 2009; Liu *et al*, 2005; Mortensen *et al*, 2011) may be partially explained by stochasticity emerging from random accumulation of proteins in autophagy deficient backgrounds over time. Similarly, difficulties with clearing cellular programs may also explain some of the reported discrepancies on the role of autophagy in reprogramming of somatic cells into pluripotent stem cells (Wu *et al*, 2015; Wang *et al*, 2013). Given enough time, autophagy deficiencies generate cellular environments in which pluripotent cells proliferate and organs are formed *de novo* without control. Once pluripotency is achieved, autophagy functions as a brake to keep subsequent tissue/organ regeneration steps at ‘cruise control’. Alltogether, our results suggest an evolutionarily conserved function of autophagy in preserving cellular homeostasis during somatic cell reprogramming. As reprogramming is central to environmental acclimation and tissue regeneration in all organisms, our findings have exciting implications for fundamental and applied biology in plants and animals.

## Materials and Methods

### Experimental model and subject details

*Arabidopsis* plants were grown in 9× 9cm pots in growth chambers at 21°C and 70% relative humidity and with an 8h photoperiod. The intensity of the light was set at 140μE/m^2^/s. The following *Arabidopsis* lines were used in this study: Columbia (Col-0), *atg2-2* (Wang *et al*, 2011), *atg5-1* (Thompson *et al*, 2005), GFP-ATG8a (Svenning *et al*, 2011) and YFP-mCherry-NBR1(Svenning *et al*, 2011). *Arabidopsis* callus was grown in 9 cm petri dishes at 21°C under continuous light. For immunoblot and proteomic experiments seedlings grown on solid MS medium (0.44% w/v agar, 1% w/v sucrose, pH5.7) were kept under 16 hours of light (150 µE/m^2^/s) at 21°C, after seed surface sterilization with 1.3% v/v bleach followed by 70% ethanol. Seedlings were then moved from solid to liquid MS media and let to acclimate for 2 days before experiments were performed.

For callus experiments, roots were excised and placed in CIM (1X Gamborg’s B5 salts with vitamins, 20 g glucose, 0.5 g/l MES, 8 g/l agar, 0.5 mg/l 2,4-D and 0.05 mg/l kinetin). The pH was adjusted to 5.7 using 1.0 M KOH. After 6 days, calli were moved to SIM (1x Gamborg’s B5 salt mixture with vitamins, 20 g/l glucose, 0.25 g/l MES, 8 g/l agar, 5 mg/l 6-(γ,γ-Dimethylallylamino)purine (2-IP), 0.15 mg/l Indole-3-acetic acid (IAA)). The pH was adjusted to 5.7 using 1.0 M KOH. For long term CIM incubation (21 days), SIM recipe was adjusted as follows: 1x MS salts with vitamins, 30g/l sucrose, 7.5 g/l agar, 0.1 mg/l IAA and 1.0 mg/l BAP. pH was set to 5.8 with 1M NaOH.

To investigate the rate of reprogramming initiation in *Physcomitrella*, the top part of 4-week-old gametophores were isolated and the tips were dissected with a surgical knife and placed on a new plate overlaid with cellophane. The gametophore tips were checked for reprogramming activation every 12 h using a Leica MZ16 F Fluorescence Stereomicroscope. For imaging, gametophore tips were dissected as described above and the individual tips were placed in small dots of 2% methyl cellulose 15cP (Sigma) in an empty Petri dish, overlaid with cellophane and cooled BCD-AT media on top (Bressendorff *et al*, 2016). Pictures were taken every 24 h using a Sony α6000 camera mounted on a Leica MZ16 F Fluorescence Stereomicroscope.

### Plant chemical treatments

For confocal microscopy, seedlings were set on liquid half strength MS media and the elongation and meristematic zone of the roots were visualized 7 days after germination. Samples were treated with half strength MS media containing one of the following: Flg22 (1 *µM*), a peptide from bacterial flagellum that elicits immune responses; PEP1 (1 µM), a small danger peptide produced upon wounding; ATP (100 µM), perceived as a danger-associated molecular pattern; ABA (1 µM), a hormone associated with abiotic stress responses; ACC (10 µM), a precursor of ethylene involved in development and senescence; BL (10 nM), a steroid hormone involved in growth; NAA (1 µM), an auxin analogue involved in growth modulation, and 6-BA (1 µM), a synthetic cytokinin regulating development.

For TSPO western blot and ABA/Flg22 proteomics experiments, plants were pre-treated with half strength MS + ABA (1 µM) for 16h and then media was replaced with half strength MS + Flg22 (1 µM) or just half strength MS for 3 h. For TSPO western blot in ABA/NAA, samples were treated as above but instead of flg22, samples were treated with MS + NAA (1 µM) for 3 hours. For Auxin/Cytokinin proteomics and catalase western blots, plants were pre-treated with half strength MS + NAA (1 µM) for 16h and then media was replaced with half strength MS + 6-BA (1 µM) or just half strength MS for 3 h. Samples were then collected, and flash frozen for further analyses.

### Confocal and light microscopy

All images were taken using a LSM700 Zeiss confocal microscope. All *Arabidopsis* root images were taken with a 63X water objective. The confocal images were analysed with Zen2012 (Zeiss) and ImageJ software.

### Protein extraction

Protein were extracted as described previously (Rodriguez *et al*, 2018). In brief, a buffer containing 50 mM Tris-HCl, pH 7.5, 150mM NaCl, 10% (v:v) glycerol (Applichem), 10 mM DTT (Applichem), 10 mM EDTA (Sigma), 0.5% (v:v), PVPP (Sigma), protease inhibitor cocktail (Roche), and 0.1% (v:v) Triton X-100 (Sigma) was used to extract proteins. Afterwards, 3x SDS with 50 mM DTT was added to the samples and this was followed by 20 min centrifugation at 4°C and 13000 x g. Supernatant was then collected and heated at 95°C for 5 min before loading samples for SDS-PAGE.

### SDS-PAGE and immunoblotting

Protein samples were separated on 12% SDS-PAGE gels, electroblotted to nitrocellulose membrane (GE Healthcare), then blocked (1 h in 5% [w:v] BSA [Merck] or 5% [w:v] milk in TBS [50mM Tris-HCl, pH 7.5 150 mM NaCl, 0.1 % -Tween-20 [Sigma]) and incubated 2 h to overnight with primary antibodies: anti-NBR1 (Agrisera) (1:5000), anti-TSPO (47), anti-GFP (TP401 AMSBio) (1:1000) or anti-CAT (Agrisera). Membranes were incubated in secondary anti-rabbit HRP conjugate (Promega; 1:5000) for 1 h. Chemiluminescent substrate (homemade or ECL Plus; Pierce) was applied before exposure to camera detection.

### Quantitative proteomics

Frozen plant material (500 mg) was lysed in lysis buffer (4% SDS, 100 mM DTT, 100 mM Tris HCl, pH7.5) and supernatant was collected after centrifugation at 20000 g for 15 min. Samples (8 µg/ml concentration) were used for mass spectrometry measurements. FASP and desalting steps were performed as previously described (Käll *et al*, 2007). These samples are then labeled with TMT according to the manufacturer’s instructions (ThermoFisher). Labelled samples were separated into fractions using an SCX system (ThermoFisher), analyzed in LC-MS/MS (Roitinger *et al*, 2015). SCX was performed using an Ultimate system (ThermoFisher Scientific) at a flow rate of 35 µl/min and a TSKgel column (ToSOH) column (5-µm particles, 1 mm i.d. x 300 mm). The flow-through was collected as a single fraction, along with the gradient fractions, which were collected every minute. In total, 130 fractions were collected and stored at −80°C.

For data analysis raw files were processed in Proteome Discoverer (version 1.4.1.14, ThermoFisher Scientific, Bremen, Germany). MS Amanda (Dorfer *et al*, 2014) (version 1.4.14.8240) was used to perform a database search against the TAIR10 database supplemented with common contaminants. Oxidation of methionine was set as dynamic modification and carbamidomethylation of cysteine as well TMT at lysine and peptide N-termini were defined as fixed modifications. Trypsin was defined as the proteolytic enzyme allowing for up to 2 missed cleavages. Mass tolerance was set to 5 ppm for precursors and 0.03 Da for fragment masses. Reporter ion intensities were extracted in Proteome Discoverer using the most confident centroid within an integration boundary of 10 ppm. Identified spectra were FDR filtered to 0.5% on PSM level using Percolator. Peptides shorter than 7 amino acids were removed from the results. Identified peptides were grouped to proteins applying strict maximum parsimony. Quantification of proteins is based on unique peptides only. Quantified proteins were exported and further processed in the R environment (version 3.4.3). Proteins were ranked by their similarity to an expected regulation pattern according to Pearson correlation. Furthermore, proteins regulated more than 1.5-fold were subdivided into clusters using k-means clustering.

### Plant propagation and high-throughput phenotyping

Seeds were stratified at 4°C in the dark for 4 days in the phytotron. The substrate (Einheitserde, ED63) was sieved (6 mm) and every single pot was filled with the same amount of substrate, 70–72 g) by using a scale to facilitate a homogenous packing density. The prepared pots were all covered with blue mats to enable a robust performance of the high-throughput image analysis algorithm (Junker *et al*, 2015). Seedlings (96) for Col-0 (WT), *atg5-1*, and *atg2-2* genotypes were propagated. The individual plants were arranged randomly by shelf. For randomization, an in-house developed R-based randomization tool was used.

The HT plant phenotyping phytotron was set to controlled conditions at 21°C during day with a night drop to 16°C at night, 60% rel. humidity and ca. 160 µmol light with a very balanced light spectrum. For the control experiment these conditions were set for the entire duration of the experiment.

For the stress experiment, the environmental conditions were changed at 21 days after sowing (DAS) to a dynamic simulation of the Swedish spring by using the hourly recorded data of the Swedish field site (Ullstorp) during 25.4. – 31.5. 2013.

The dry weight was scored by harvesting a random sample of 36 replicates per genotype (plants cut off at the base just above the soil).

This particular phytotron allows high-throughput plant phenotyping with the integrated and automated x,y,z sensor-to-plant RGB imaging system, delivering images of 1260 plants in one go. Images were taken twice a day during standard conditions; in the morning 1 h after the lights went on and in the late afternoon, 1 h before lights went off. During the Swedish conditions the number of pictures per day was increased to 5 in order to score possibly quick changes in the phenotypes. In total, 102,059 pictures were taken for the experiment with the simulation of Swedish spring.

For the data processing, quality control and preliminary analysis we have used PHENOComp, an R package developed by the VBCF BioComp facility. The R programming environment (R Core Team, 2018) was used for performing the statistical analysis and generating the visualizations for the Figs. Data clean-up and transformations were done using the tidyr package (https://cran.r-project.org/package=tidyr). Datasets were checked for normality and heteroscedasticity to select the appropriate statistical tests. Statistical significance for differences in mean values for dry weight and seed weight was determined using the Kruskal-Wallis non-parametric test. Statistical significance for differences in the variance was determined using Levene’s test from the car package (https://cran.rproject.org/package=car). In both cases, pairwise comparisons were calculated using Dunn’s test as implemented in the PMCRM package (https://cran.r-project.org/package=PMCMR) and corrected for multiple testing using the Holm method. The post-hoc test for pairwise comparisons of the variances between groups was calculated using the absolute values of the residuals (deviation from the median of the group), as recommended in (Boos & Brownie, 2005). We tested for difference in the coefficient of variation using the asymptotic Feltz and Miller test for the equality of coefficients of variation (Feltz & Miller, 1996), as implemented in the package cvequality (https://cran.rproject.org/package=cvequality). Plots were produced using the packages ggplot2 (https://CRAN.R-project.org/package=ggplot2) and ggpubr (https://CRAN.Rproject.org/package=ggpubr.”

## Acknowledgments

We would like to thank Henri Batoko (UCL, Louvain-la-Neuve) for anti-TSPO antibody. Anne Simmonsen (IBMS, Oslo), Daniel Klionsky (U.Michigan) and John Mundy (U.Copenhangen) for critical reading of the manuscript. Rune Salomonsen for technical assistance. Microscopy was performed at the Center for Advanced Bioimaging, University of Copenhagen.

## Funding

Work was supported by grants to: M.P.:Novo Nordisk Fonden (NNF16OC0021618 2017); Y.D.: Austrian Academy of Sciences and WWTF (Project No: LS17-047).; K.M. Austrian Science Fund (SFB F3402, TRP 308-N15) and ERA-CAPS (I 3686); P.S.D.: Interreg-RIATCZ project.

## Author Contribution

Conceptualization: M.P. and E.R.; Investigation: E.R., J.C., J.A., V.K., J.O., J.C., C.L. M.S., S.K., J.J. and Z.Z; Formal analysis: E.R., J.C., J.A., J.O., G.D., K.M., P.S.D., Y.D.; Resources: S.S, S.L.; Writing: M.P., Y.D. and E.R. Visualization: E.R., J.C., Y.D.; Supervision: M.P. Y.D, E.R.; Funding Acquisition M.P. and Y.D. All authors read the manuscript and agreed with the findings reported.

## Supplementary Figures

**Supplementary Fig.1.**
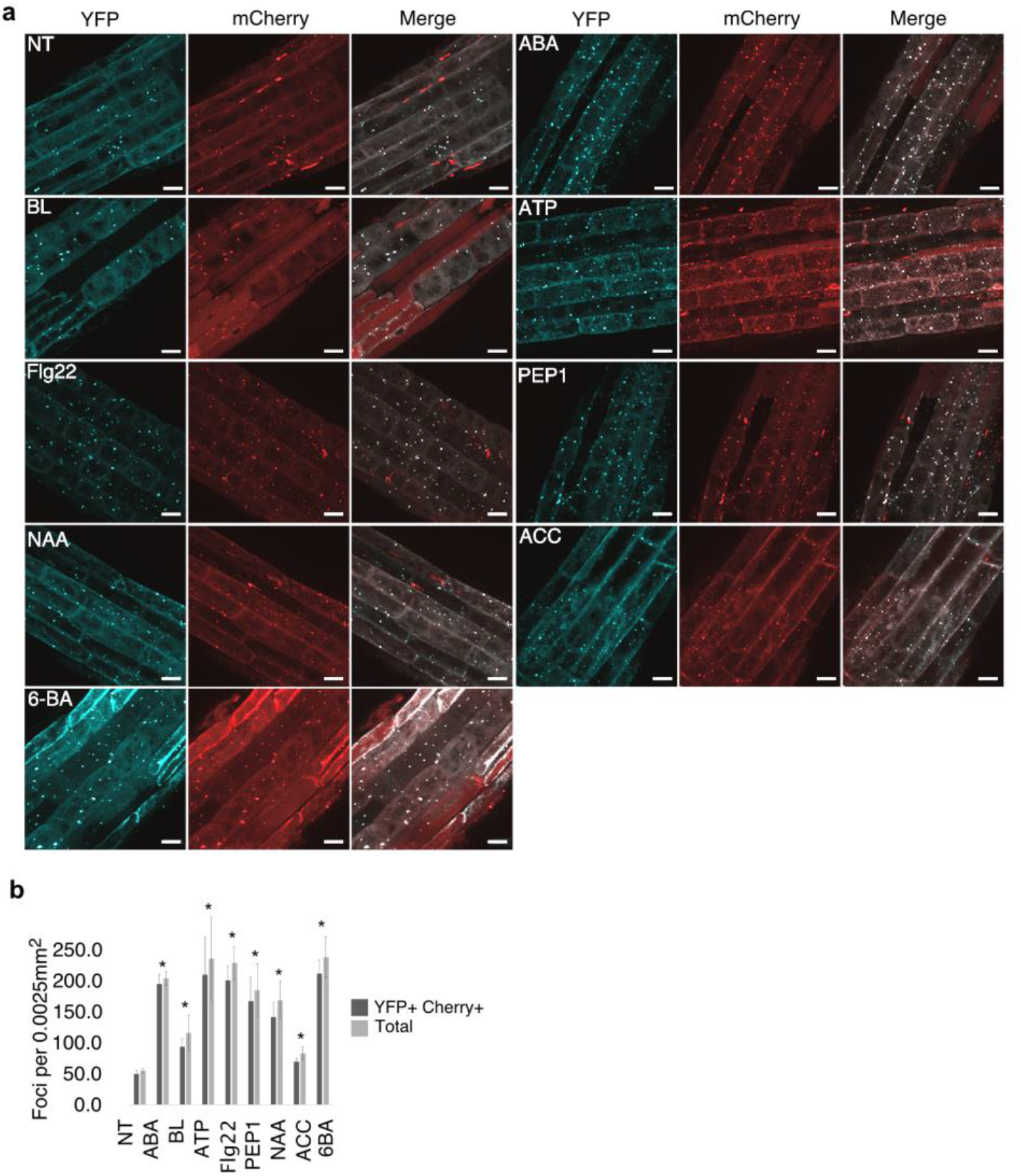
Autophagy is rapidly induced upon recognition of a wide range of stimuli. YFP-mCherry NBR1 accumulation before or 30 min after treatment with ACC, ABA, ATP, BL, 6-BA, Flg22, NAA and PEP1. (**a**) Representative maximum intensity projection images of 10 Z-stacks per image. (**b**) Quantification of YFP/mCherry foci for given treatments per 0.0025 mm^2^. Values are based on 3 independent experiments, with 3 individuals per condition. Bars marked with an asterisk (*) are statistically significant (P<0.05).

**Supplementary Fig. 2.**
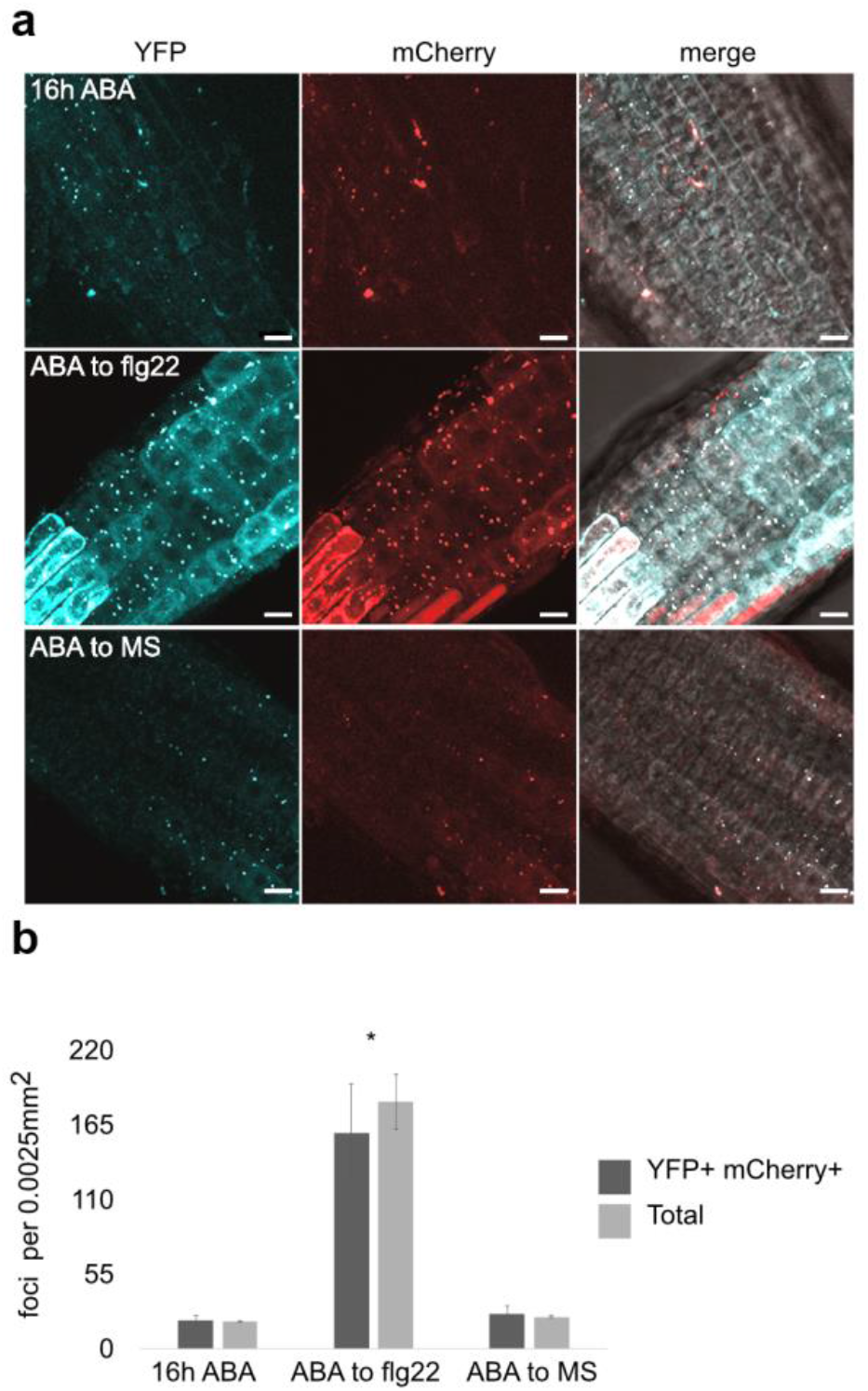
Autophagy is reactivated upon contrasting stimuli perception. Seedlings were acclimated for 16 h in MS containing ABA and then imaged 30 min after being swapped to MS or MS containing flg22. (**a**) Representative maximum intensity projection of 10 Z stacks per image. Experiments were repeated, independently, 3 times with similar results Scale bar: 10 µm. (**b**) Quantification of YFP/mCherry foci for given treatments per 0.0025 mm^2^. Values are based on 3 independent experiments, with 3 individuals per condition. Bars marked with an asterisk (*) are statistically significant (P<0.05).

**Supplementary Fig. 3.**
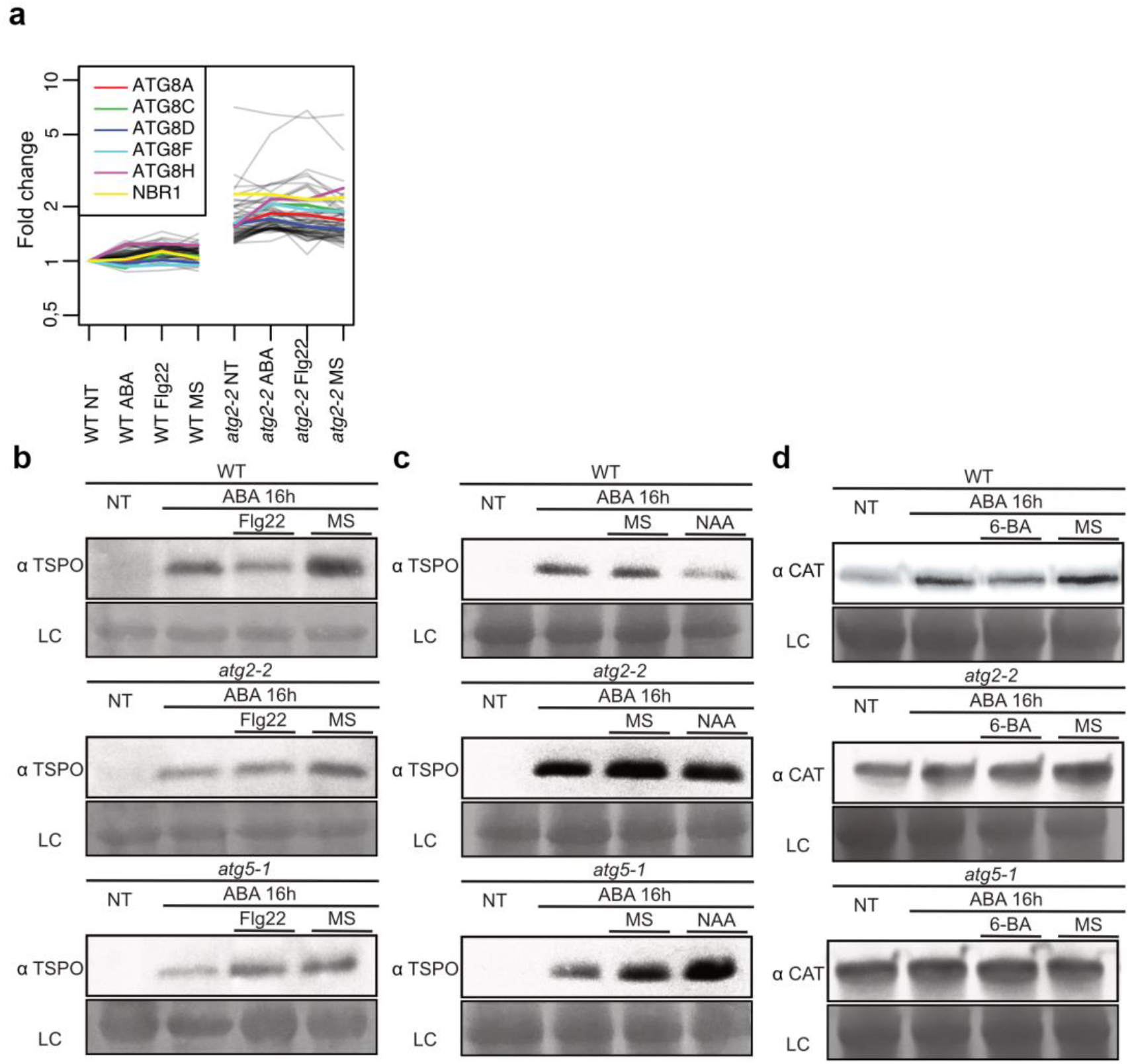
Protein level change upon treatments with consecutive stresses. (**a**) Proteins that accumulate in *atg2* in comparison to WT. (**b**) TSPO immunoblot for WT, *atg2-2* and *atg5-1* samples treated as described in Fig. 2A for the ABA to flg22 consecutive stress set. (**c**) TSPO immunoblot for WT, *atg2-2* and *atg5-1* samples treated as described in Fig. 2A for the ABA to NAA consecutive stress set. (**d**) CAT immunoblot for WT, *atg2-2* and *atg5-1* samples treated as described in Fig.2A for the NAA to 6-BA consecutive stress set.

**Supplementary Fig. 4.**
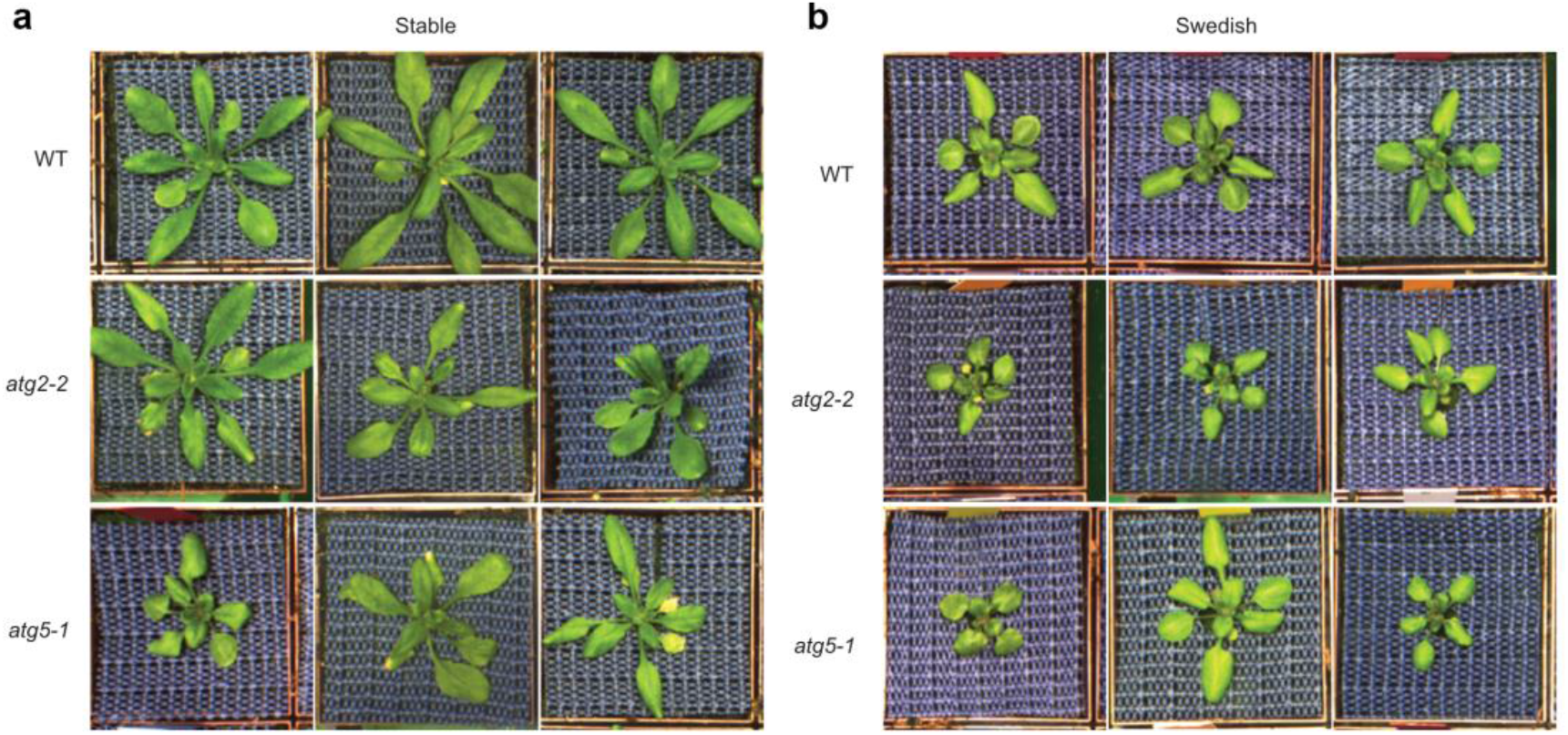
Representative picture of WT *atg2-2* and *atg5-1* grown in stable or Swedish spring 2013 conditions. (**A**) WT, *atg2-2* and *atg5-1* grown in stable conditions. (**B**) WT, *atg2-2* and *atg5-1* grown in Swedish spring 2013 conditions.

**Supplementary Fig 5.**
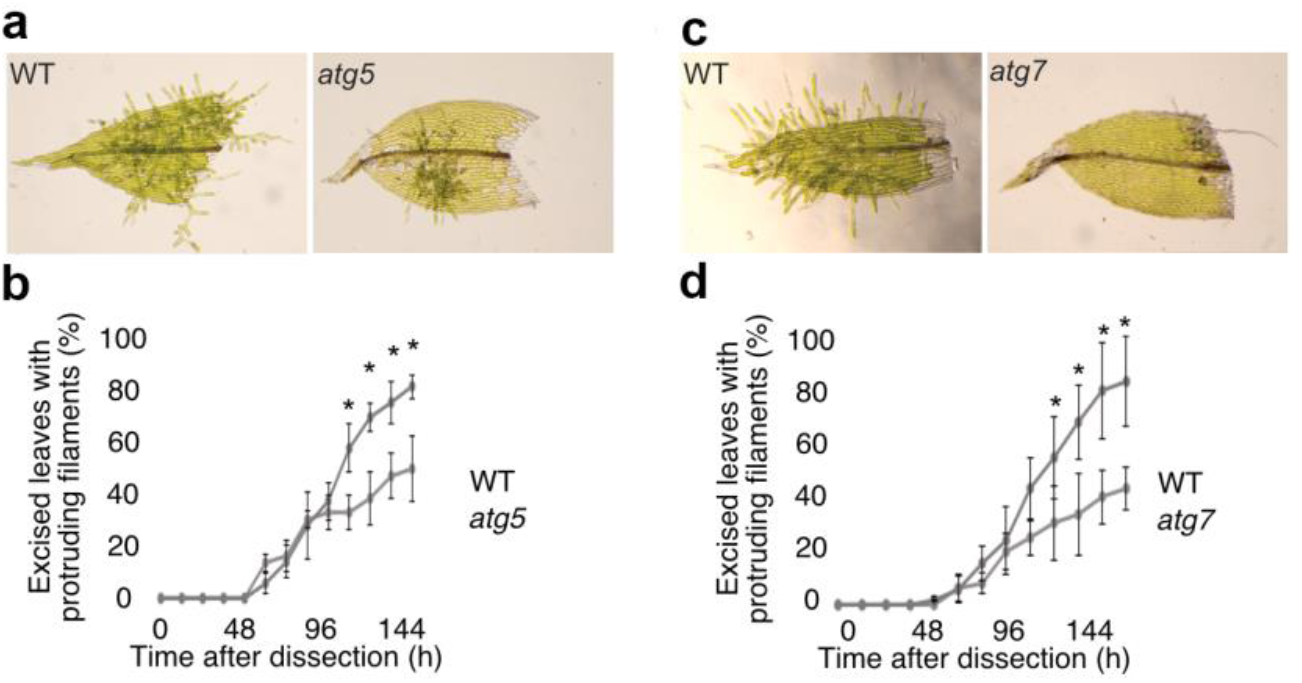
Autophagy is necessary for wound-induced dedifferentiation and tissue repair in *Physcomitrella patens*. (**A**) Representative Fig. of gametophore leaves from WT (Gransden) and *atg5*, undergoing dedifferentiation and cell protrusion 144h after wounding. (**B**) Number of gametophore leaves displaying cell protrusions after wounding for the genotypes given. (**C**) Representative Fig. of gametophore leaves from WT (Reuter) and *atg7*, undergoing dedifferentiation and cell protrusion 144h after wounding. (**D**) Number of gametophore leaves displaying cell protrusions after wounding for the genotypes given. Results are given as mean plus standard deviation of the mean of at least 30 individuals from 3 independent experiments.

**Supplementary Fig. 6.**
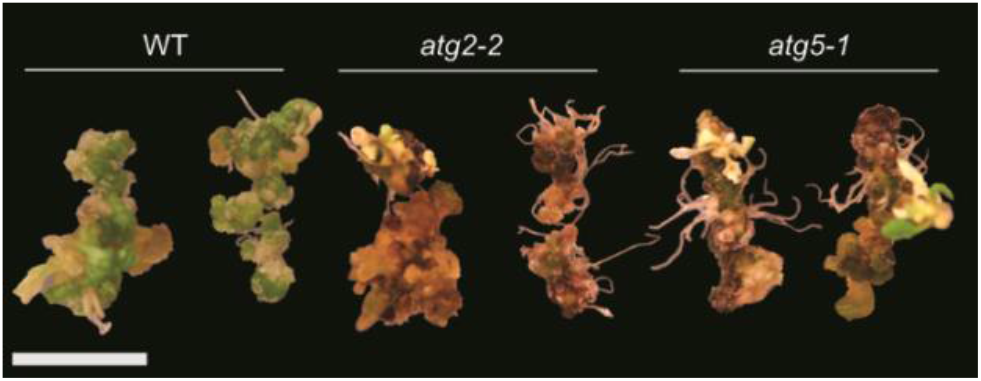
Calli derived from autophagy mutants senesence prematurely and die after prolonged growth on SIM. Representative picture of WT, *atg2-2* and *atg5-1* calli grown on CIM for 21 days and then kept on SIM for 35 days.

## Supplementary Tables

**Table S1**. Proteins that differentially accumulate upon treatment following the experimental setup represented in Fig. 2A for ABA to flg22 set up.

**Table S2**. Proteins that differentially accumulate upon treatment following the experimental setup represented in Fig. 2A for NAA to 6-BA set up.

